# Fasting increases investment in soma upon refeeding at the cost of gamete quality in zebrafish

**DOI:** 10.1101/2022.07.18.500454

**Authors:** Edward R Ivimey-Cook, David S. Murray, Jean-Charles de Coriolis, Nathan Edden, Simone Immler, Alexei A. Maklakov

## Abstract

Fasting increases lifespan in invertebrates, improves biomarkers of health in vertebrates, and is increasingly proposed as a promising route to improve human health. Nevertheless, little is known about how fasted animals use resources upon refeeding, and how such decisions affect putative trade-offs between somatic growth and repair, reproduction, and gamete quality. Such fasting-induced trade-offs are based on strong theoretical foundations and have been recently discovered in invertebrates, but the data on vertebrates is lacking. Here we report that fasted female zebrafish, *Danio rerio*, increase investment in soma upon refeeding but it comes at a cost of egg quality.

Specifically, an increase in fin re-growth was accompanied by a reduction in 24-hours post-fertilization offspring survival. Refed males showed a reduction in sperm velocity and impaired 24-hour post-fertilisation offspring survival. These findings underscore the necessity of considering the impact on eggs and sperm when assessing evolutionary and biomedical implications of lifespan-extending treatments in females and males and call for careful evaluation of the effects of fasting (both during and post) on fertilisation.

## Introduction

In the past two decades there has been a wealth of research highlighting the robust lifespan- and healthspan-extending effects of dietary restriction (DR) [1–3], defined as a reduction in nutrient intake without malnutrition. Much of this work has been directed towards the trade-off between reproduction and survival that typically occurs as part of an organism’s DR response. Moreover, there are the two predominant evolutionary theories which seek to explain why this lifespan extension occurs [4,5]. Firstly, the resource allocation theory [5] suggests that organisms should reallocate resources away from reproduction into somatic maintenance and repair under periods of food limitation. In doing so, there is a negative effect on reproduction but an increase in the likelihood of surviving until food becomes plentiful again [5]. A more recent theory, suggests that this lifespan extension is merely a by-product of upregulating autophagy and the breakdown of internal resource (in addition to a general laboratory artefact of reduced selection pressures) in order to maximise fitness in dietary restricted environments [4]. However, regardless of these two theories, which differ in the ultimate and proximate reasons for DR-mediated lifespan extension, there is a general observation across multiple taxa that increased survival during periods of transient fasting is often associated with reduced reproduction.

More recently, the adaptive nature of the DR response has come under scrutiny [6,7], particularly when individuals are brought back into an *ad libitum* environment. Whereas reduced reproduction during fasting could be adaptive when resources are limited, reducing the likelihood of producing offspring into a resource-poor environment and increasing the likelihood of surviving until the environment becomes resource-rich again, there is uncertainty about the effects of DR on survival and reproduction upon refeeding. Indeed, recent work in *Drosophila melanogaster* has shown that upon exiting a period of DR, not only is there an increase in mortality [6–8] but, under some circumstances, there is no compensation for this survival decline with increased fecundity, suggesting an overall cost of the DR response [5]. However, more recently work by Sultanova et al. (2021) has shown a contrasting effect, with increased fecundity and mating behaviour in post-DR flies who suffered a similar increase in mortality [6]. There is therefore, a distinct lack of consensus on whether the DR response is associated with increased reproduction and fitness upon refeeding, but there is also a lack of data on vertebrate species that extends beyond measuring the physiological effects of refeeding, including weight gain and quantity of adipose tissue ([8,9] although see [10]).

Another important factor to consider when determining the fitness effects of dietary restriction is the inter-(from parent-to-offspring) or trans-generational (from parents-to-grand-offspring-and-onwards) effects manifesting on future generations. In particular, recent studies in *Caenorhabditis elegans* and *C. remanei* [12–16] have revealed the significant impact that parental fasting can have on various measures of offspring performance, from mediating egg size and number [12] to influencing future survival in both immediate and distant generations [13]. Nevertheless, the effects of DR on fitness of offspring produced after parents resumed their normal diet is poorly understood and effects could be reversed if post-DR parents are in a better physiological condition. For instance, in a natural population of female Brandt’s vole (*Lasiopodomys brandtii*) refeeding after a period of 70% food restriction caused survival rate of weaning offspring to increase by ~10% suggesting that individuals were able to overcompensate for the reduced reproduction during dietary restricted period. However, it has been widely recognised that both maternal (egg- or care-mediated) and paternal (sperm- or care-mediated) phenotypes can readily and separately contribute to changes in offspring condition [17–25]. Thereby not only does this warrant more research into other vertebrate taxonomic groups, but also to discuss the effects of refeeding considering both maternal and paternal effects. This is especially relevant as the effects of DR have been found to differ markedly depending on the sex of the experimental individual (see [3]).

To this end, we investigated the effects of dietary restriction and subsequent refeeding on various important life history characteristics in the model vertebrate system, the zebrafish (*Danio rerio*). We focused on addressing three key questions 1) How do individuals allocate resources upon refeeding between somatic maintenance, fecundity, and offspring quality? 2) Do these responses differ between females and males? 3) Do periods of fasting and refeeding have differing effects on offspring lifehistory traits such as survival and growth? We found the female zebrafish invest in their somatic maintenance upon refeeding and are better able to deal with somatic injury compared to control counterparts. However, this benefit comes at a cost of lower egg quality. Furthermore, we found that DR reduces quality of male gametes when males resume their normal feeding. Our findings highlight the importance of considering all aspects of an individual’s life history from fertilisation and gametes to somatic maintenance and repair, in order to fully understand the implications of transient dietary restriction.

## Methods

### Animal husbandry and diets

All experimental assays used zebrafish (*Danio rerio*) obtained from a population of outbred wild-type (AB strain) that originated from the European Zebrafish Research Centre (EZRC, Tübingen, Germany), raised to sexual maturity (≥ 6months old) and were maintained for up to two generations following an outbreeding regime at the Controlled Ecology Facility (CEF) at the University of East Anglia (UEA, UK). Individual fish were housed in three litre tanks in a recirculating rack system (ZebTec Active Blue Techniplast UK Ltd) at 26.4±1.4°C and kept on a 12L:12D light cycle. When fed *ad libitum* and prior to the experimental assays, they were fed two or three times a day with a mixture of live artemia (Brineshrimp cysts, ~100 artemia per ml of water) and dry food (400-600μm, Sparos Zebrafeed) and kept at 1:1 sex ratios. Individuals were then placed into one of two diets for 15 days, either fully-fed (*ad libitum*) or fasted (no food). After 15 days, the fasted individuals were placed back onto *ad libitum* food. Lastly, as zebrafish can display social dominance traits, leading to aggression against subordinate individuals; artificial aquaria plants were added for sheltering and hiding space to prevent aggressive interactions from influencing results within the current study [26].

Experimental protocols were approved by the Named Animal Care & Welfare Officer (NACWO) (project licence number: P0C37E901).

### Caudal fin assay

To measure somatic growth, on day three of the experimental treatment a section (≥50%) of the caudal fin of 24 fed and 36 fasted fish (12 fed and 18 fasted males and females spread over five replicate blocks) was removed. The caudal fin, beginning from the caudal peduncle to the tip of each tail lobe, was then photographed, including a scale bar, and the surface area measured using ImageJ software [27] before and immediately after removal, and then on days 7, 15, 21, 28 and 35.

### Reproductive assay

To test various measures of reproductive performance, 30 experimental males and 30 experimental females (again, with 12 fed and 18 fasted males and females) were paired on days 7, 15, 21, 28 and 35 with wildtype fish of the opposite sex from the general stock population and placed in a breeding tank separated by a divider to allow for odour acclimatisation (1 litre Breeding Tanks, Techniplast). On the morning of breeding, the divider was removed and pairs were checked every hour to monitor spawning success. Fish were given a maximum of five hours to spawn, upon which, if they were unsuccessful, were given another wild-type partner and the process repeated the next day. Upon spawning, eggs were transferred into a Petri dish with tank water and Methylene Blue (0.1%) to prevent fungal infection. The eggs were then assessed after 2 and 24 hours. Eggs were compared to embryonic diagrams produced by Kimmel et al. (1995), which describe stages of zebrafish embryonic development. Eggs which deviated from the diagrams were classified as abnormal Eggs were classified as dead if they did not contain a yolk or if they were dark green or black in colour, or unfertilised if there was no cell division within the egg 2 hours after sperm had been added [28]. Lastly, eggs were taken from day 5 of reproduction and used in the subsequent growth assay. Here, fry length was measured at three and five days post-fertilisation. In total, fry from 526 fed and 400 fasted individuals were measured (N.B. the total number was higher, but some were omitted as they were deformed or the eggs did not hatch; in addition, several individuals (~124) only provided estimates of growth on one day, these were omitted from the additional analysis of total growth).

### Sperm assay

Ejaculates were collected from each experimental male (12 for fasted and 12 for control fed) on days 7, 15, 21 and 35. To do this, all experimental fish were transferred to one litre breeding boxes which contained an internally perforated tank. Fish from the same tank were kept together and were kept in absolute darkness at 28°C for 17 hours without food to avoid contamination of samples with faeces. The males were then anesthetized in in 10mg L^-1^ AquaClam™ (Western Chemicals Inc.) solution for a maximum of 60 seconds, briefly rinsed in system water and placed ventral side up into a moist sponge under a stereomicroscope. A paper towel was used to dry the area immediately adjacent to the ventral pore to avoid sperm coming into contact with water and prematurely activating. Sperm was then released by gently pressing the belly and collected in a micro capillary tube (Sigma-Aldrich, P0674). Collected sperm was added to microtubes containing 20μl premix solution, consisting of 1:9 Hank’s buffer to dH20 and placed on ice. Each sample was analysed within two hours of extraction using a Computer Assisted Sperm Analysis (CASA) system. For each sample, 2μl of ejaculate-premix was activated with 3μl of tank water. Of this 5μl total mix, 3μl were transferred to a chambered Cytonix MicroTool B4 Slide. Sperm movement was recorded at 100x magnification using a 782M monochrome CCD progressive camera and a UOPUB203i trinocular microscope. The parameter settings used during this assay with the ISAS v. 1 software were as follows: 50 fps, 50 frames used, 5-50μm^2^ particle area. We primarily assess curvilinear velocity (VCL) defined as 10-45μm/s for slow, 45-100μm/s for medium and rapid as >100μm/s.

### Statistical analysis

All analyses were conducted using R v. 4.1 [29]. Data was tidied and cleaned using the tidyverse v. 1.3.1 [30], hablar v. 0.3.0 [31] and janitor v. 2.1.0 [32] collection of packages. Linear mixed effect models were run using glmmTMB v. 1.1.2.3 [33,34] and checked for zero-inflation, dispersion and overall fit using DHARMa v. 0.4.5 [35]. Estimated marginal means for each model were then created using emmeans/emtrends v. 1.7.2 [36] and visualised using the ggeffects v. 1.1.1 package [37]. Lastly, overall significance was tested using type 3 anovas from the car v. 3.0-12 package [38].

For each trait, aside from curvilinear velocity (VCL) and offspring survival, we ran two models. The first contained data from both males and females to test for differences in (parental)-sex-specific responses. The second (and third) were female- and male-specific models which looked more closely at the differences between *ad libitum* and fasting treatments. Lastly, for all traits aside from fry growth, the analysis was split into *during* and *post*-fasting (or refeeding) in order to observe any differences in life history trade-offs between these two dietary events.

Models for all traits were fit as follows. For fin regrowth and fry growth per day, dietary treatment (twoway factor: fed/fasted) was fit in a Gaussian model and included an interaction with the linear form of day (and if analysing both sexes together, with a three-way interaction of sex; note that in all cases sex was referring to the experimental individual in the parental generation) and the random effect of fish id nested within experimental replicate. For average hourly fry growth, a similar model to above was fit but due to model convergence issues when considering the two-way interaction between treatment and parental sex (day was no longer included), replicate was instead fit as a fixed effect and the model was run without a random effect structure. Reproduction was measured in two distinct ways. Firstly, age specific reproduction was fit with a similar structure to above (either a three-way interaction of the linear form of time, with treatment, and sex or if in the sex-specific model with simply a two-way interaction between treatment and time), albeit with a negative binomial error distribution (treatment interacting with day, random effects of fish id nested within experimental replicate). Secondly, lifetime reproduction success (LRS), or the total number of offspring produced either in the fasting/refeeding period, was fit in a negative binomial model (in both cases, a Poisson model was originally considered but rejected due to poor fit and significant overdispersion) with simply treatment as the main fixed factor (aside from the combined sex model, which again, included a two-way factor of sex) and a random intercept of experimental replicate. In both cases, if significant zero-inflation was detected in the residuals of model (identified using DHARMa) then an additional zero-inflated component was added to each model. Offspring survival was modelled as a binomial response (survive = 1, dead = 0) where dietary treatment interacted with day and the same random effect structure as age-specific reproduction was used. Lastly, log-transformed sperm VCL was analysed in a gaussian model with the same fixed effect structure (treatment:day) however a two-level factor of replicate block (two level factor) was also added. Only fish id was included as a random effect.

## Results

See Table S1 for a full overview of the sex-specific results of fasting and refeeding.

### Effect of transient fasting on Soma

When assessing regrowth during the fasting period, there were no significant differences over time between males and females that were fasted or fed (X^2^ = 1.27, *p* = 0.260). However, despite the lack of statistical significance, fasted males and females had increased regrowth, albeit this effect was more pronounced in males (female fed: 0.37 - female fasted: 0.39, estimate = −0.02 (0.05), *p* = 0.748; male fed: 0.33 - male fasted: 0.43, estimate = −0.10 (0.052), *p* = 0.057; Table S2). Similarly, during the period of refeeding, males and females did not differ in regrowth rates between the fasted and fed treatments (X^2^ = 0.71, *p* = 0.400). However, across both sexes, fasted individuals had increased regrowth in comparison to their fully-fed counterparts (fully-fed: 0.22 - fasted: 0.27, estimate = −0.048 (0.02), *p* = 0.020; Fig. 1; Table S3). This significance was largely driven by post-fasting females, and not males, as they exhibited higher fin regrowth in comparison to those that were fed *ad libitum* (female fed: 0.16 - female fasted: 0.36, estimate = −0.20 (0.06), *p* = 0.001; Fig. 1; Table S4).

**Figure 1.**
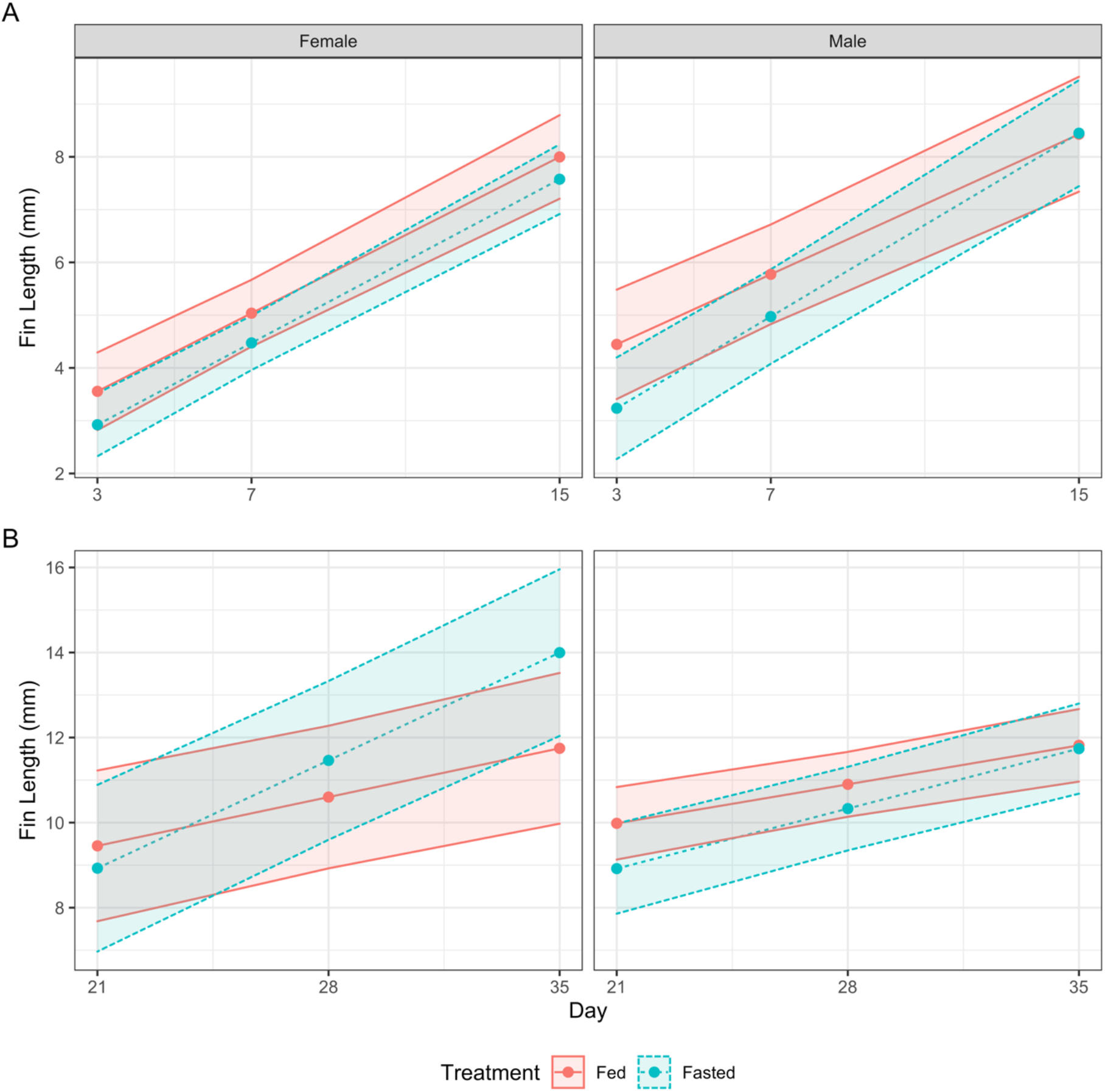
Sex-specific caudal fin regrowth prior to refeeding (A) and after refeeding (B). Colours represent fasted (blue, dashed) or fed (red, solid) dietary treatments in females (left) or males (right). Lines, points and 95% confidence intervals represent predicted values from a linear model.

### Effect of transient fasting on offspring quality and quantity

Transient fasting had sex-specific effects on the reproduction of males and females. When looking at both sexes combined, there were no differences in age-specific reproduction during the period of fasting (X^2^ = 0.66, *p* = 0.416). Only in females was the average reproduction across both days 7 and 15 significantly lower in fasted individuals (EMM fed females: 76.8, fasted females: 40.1, contrast = 1.92, *p* = 0.0169; Table S5). This resulted in a reduced LRS for females but not males during the dietary treatment period (LRS: Female fed = 138.1, female fasted = 63.7, estimate = 2.17, *p* = 0.030; male fed = 175, male fasted = 162, estimate = 1.08, *p* = 0.726; Table S6-S7). This difference between fasted and fed individuals disappeared when females began to refeed (average reproduction EMM fed females: 88.4, fasted females: 100.1, contrast = 0.88, *p* = 0.566; Table S8) resulting in comparable LRS between fed and fasted treatments (although we note the large increase of LRS for females as they transition from fasting to refeeding; LRS: fed = 240, fasted = 281, estimate = 0.856, *p* = 0.509; Table S9). Male LRS on the other hand was unaffected by both the period of fasting (see above) and refeeding, with fasted individuals showing reduced reproduction (albeit not significantly so – LRS EMM Fed males: 304.0, Fasted males: 218.0, estimate = 1.40, *p* = 0.162; Table S9). This reduced reproduction during fasting coincided with the relatively high egg survival of fasted females in comparison to fed females, although we note that fasted females still showed a significant decline (EMM fed females = 0.046, fasted females = −0.072, estimate = 0.12, *p* = 0.009; Fig. 2; Table S10). In contrast, males despite not showing a detectable decline in offspring number, produced a large negative effect of fasting on the egg survival of their mate (EMM fed males = −0.01, fasted males = −0.47, estimate = 0.46, *p* <0.001; Fig. 2; Table S11). Upon refeeding, the difference in trend between fasted and fed males became smaller largely owing to a reduction in the negative effects of fasting, however the effect size between the two treatments was still statistically different (EMM fed males = 0.017, fasted males = −0.15, estimate = 0.17, *p* <0.001; Fig. 2; Table S12). For females, the opposite occurred, with fasted females seemingly trading off increased offspring number with a greater reduction in egg survival when fed *ad libitum* (EMM fed females = 0.11, fasted females = −0.14, estimate = 0.25, *p* <0.001, denoted by the further decrease in slope value between fasting and refeeding periods; Fig. 3; Table S13).

**Figure 2.**
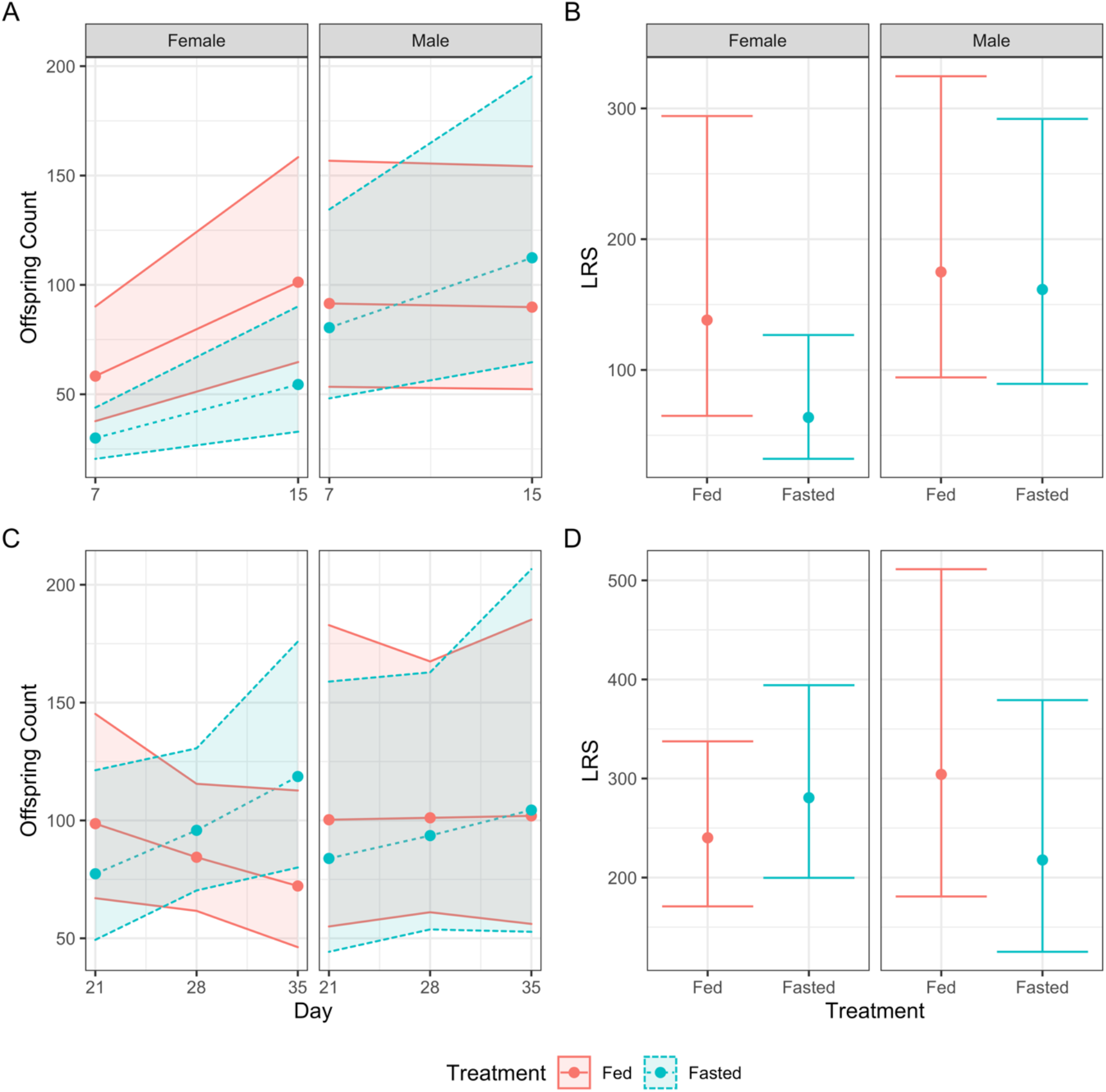
A & C) Age-specific reproduction. Colours represent fasted (blue) or fed (red) dietary treatments in females (left) or males (right). Lines, points and 95% confidence intervals represent predicted values from a linear model. B & D) Lifetime reproduction success (LRS) in either female (left, red) or males (right, blue). Lines, points and 95% confidence intervals represent predicted values from a linear model. A & B represent reproduction whilst individuals are within their respective dietary treatments. C & D are when all individuals have been placed back into *ad libitum*.

**Figure 3.**
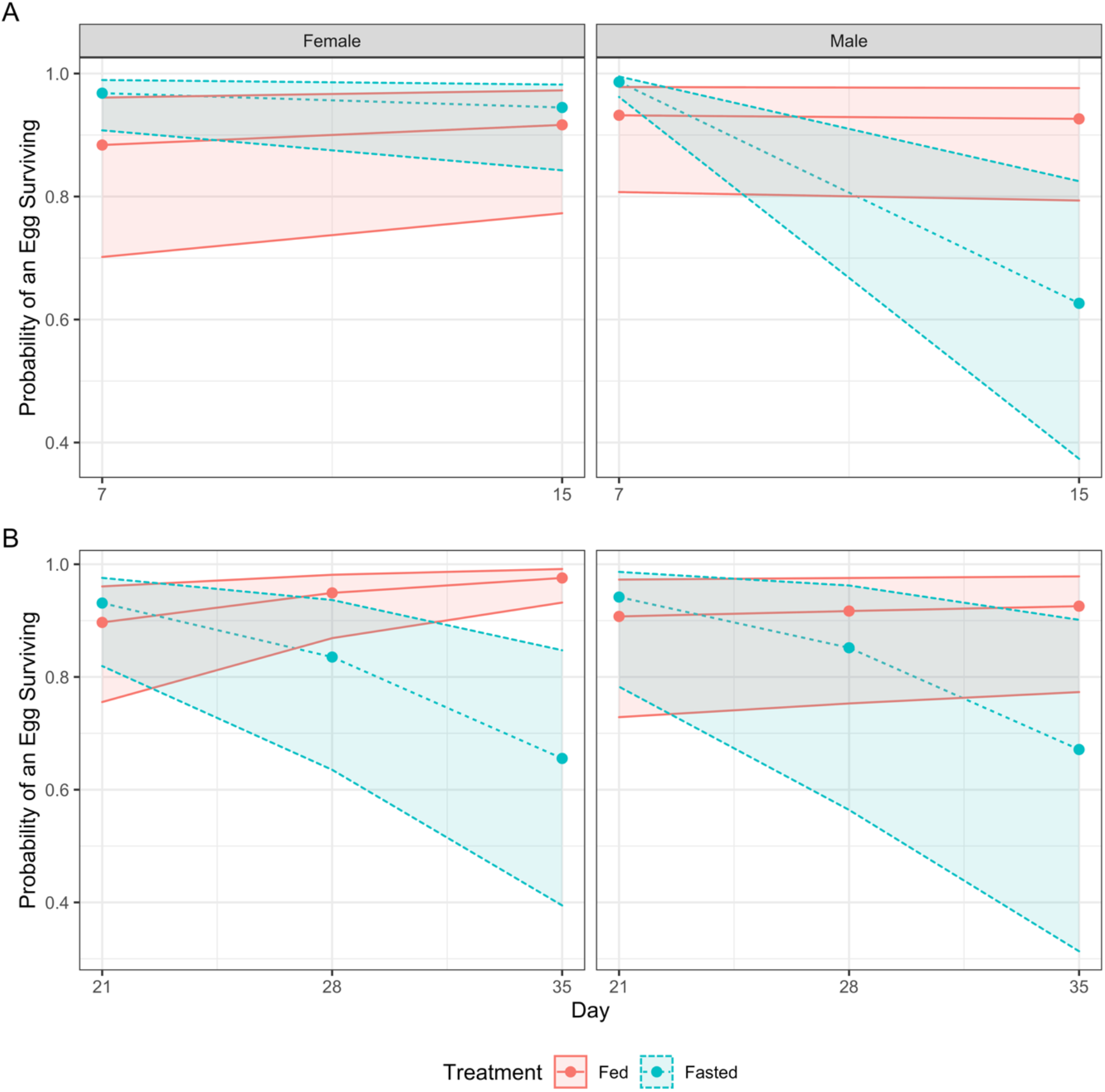
Age-specific egg survival during-(A) or post-fasting (B). Colours represent fasted (blue) or fed (red) dietary treatments in females (left) or males (right). Lines, points and 95% confidence intervals represent predicted values from a linear model.

### Effect of transient fasting on sperm quality

Male sperm quality, assessed through velocity curvilinear (VCL) declined with time during both the fasting and refeeding periods (Fig. 6A & B). However, during the fasting period, this decline was greater for fed individuals (VCL EMM: Fed = −0.009, Fasted = −0.003, estimate = −0.006, *p* = 0.001; Fig. 4; Table S14). The opposite was true during the period of refeeding, the fasted individuals exhibited a faster decline in VCL with time (EMM: Fed = −0.008, Fasted = −0.018, estimate = −0.01, *p* <0.001; Fig. 4; Table S15).

**Figure 4.**
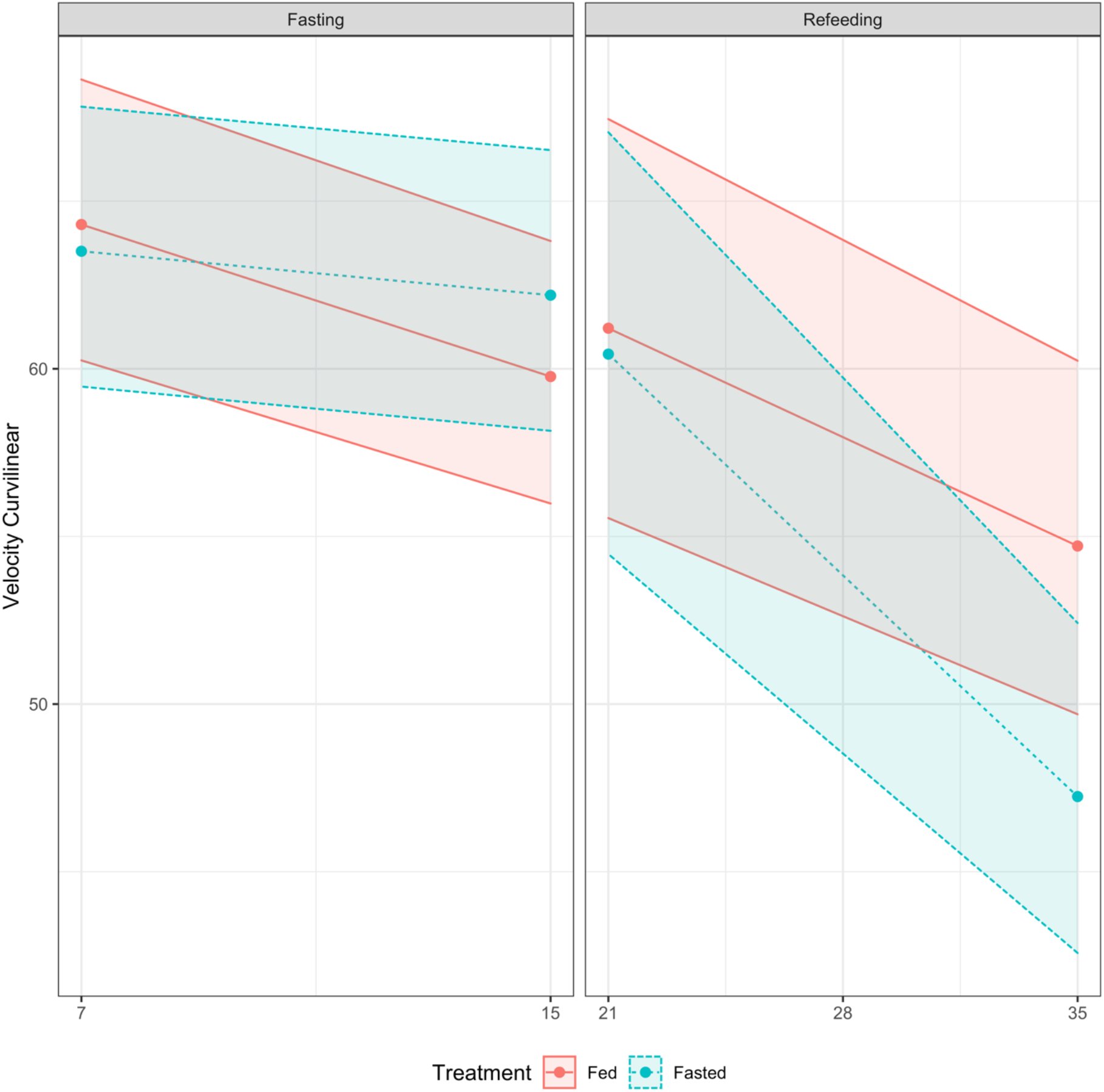
Age-specific sperm velocity curvilinear (VCL) during-(left) or post-fasting (right). Colours represent fasted (blue) or fed (red) dietary treatments in males. Lines, points and 95% confidence intervals represent predicted values from a linear model.

### Effect of transient fasting on fry growth

Sexes did not differ in average fry growth per day, with offspring from both fasted and fed males and females having similar growth rates (X^2^ = 0.161, *p* = 0.689). However, across both treatments, offspring from males had faster average growth per day than offspring from females (Growth rate EMM: Females = 0.28, Males = 0.33, estimate = −0.05, *p* <0.001; Fig. 5; Table S16). In a similar manner, across both sexes, offspring from fasted individuals grew faster than those from *ad libitum* parents (EMM: Fed = 0.29, Fasted = 0.32, estimate = −0.03, *p* = 0.025; Fig. 5; Table S17). When looking within each sex, growth rates for offspring from fasted and fed males and females were not different from each other (EMM: Fed Males = 0.31, Fasted Males = 0.34, estimate = −0.03, *p* = 0.148; Fed Females = 0.26, Fasted Females = 0.30, estimate = −0.04 *p* = 0.074; Fig. 5; Table S17-18). although we note that in both cases offspring from fasted individuals had faster growth rates, albeit not statistically significant. This difference however, may have been driven in part by unequal sampling between time points on days three and five. In light of this, we refocused the analysis to investigate average growth rates per hour (Fig. 5) in order to obtain a higher resolution of analysis. Offspring from both males and females did not differ in their growth rates between the fed and fasted parental treatments (X^2^ = 0.01, *p* = 0.911). However, across both parental treatments, offspring from male individuals had increased per hour growth rate (EMM: Female = 0.012, Male= 0.014, estimate = −0.002, *p* = 0.023; Fig. 5; Table S19). Similarly, offspring from fasted individuals across both sexes had increased growth (EMM: Fed = 0.012, Fasted = 0.014, estimate = −0.002, *p* = 0.010; Fig. 5; Table S19). This difference was largely driven by offspring from male individuals, where in the sex-specific analysis only the difference between the hourly growth of the offspring from fed males and fasted males were significant (EMM: Male Fed = 0.013, Male Fasted = 0.015, estimate = −0.002, *p* = 0.028; Female Fed = 0.011, Female Fasted = 0.012, estimate = −0.001, *p* = 0.146; Fig. 5; Table S20-S21).

**Figure 5.**
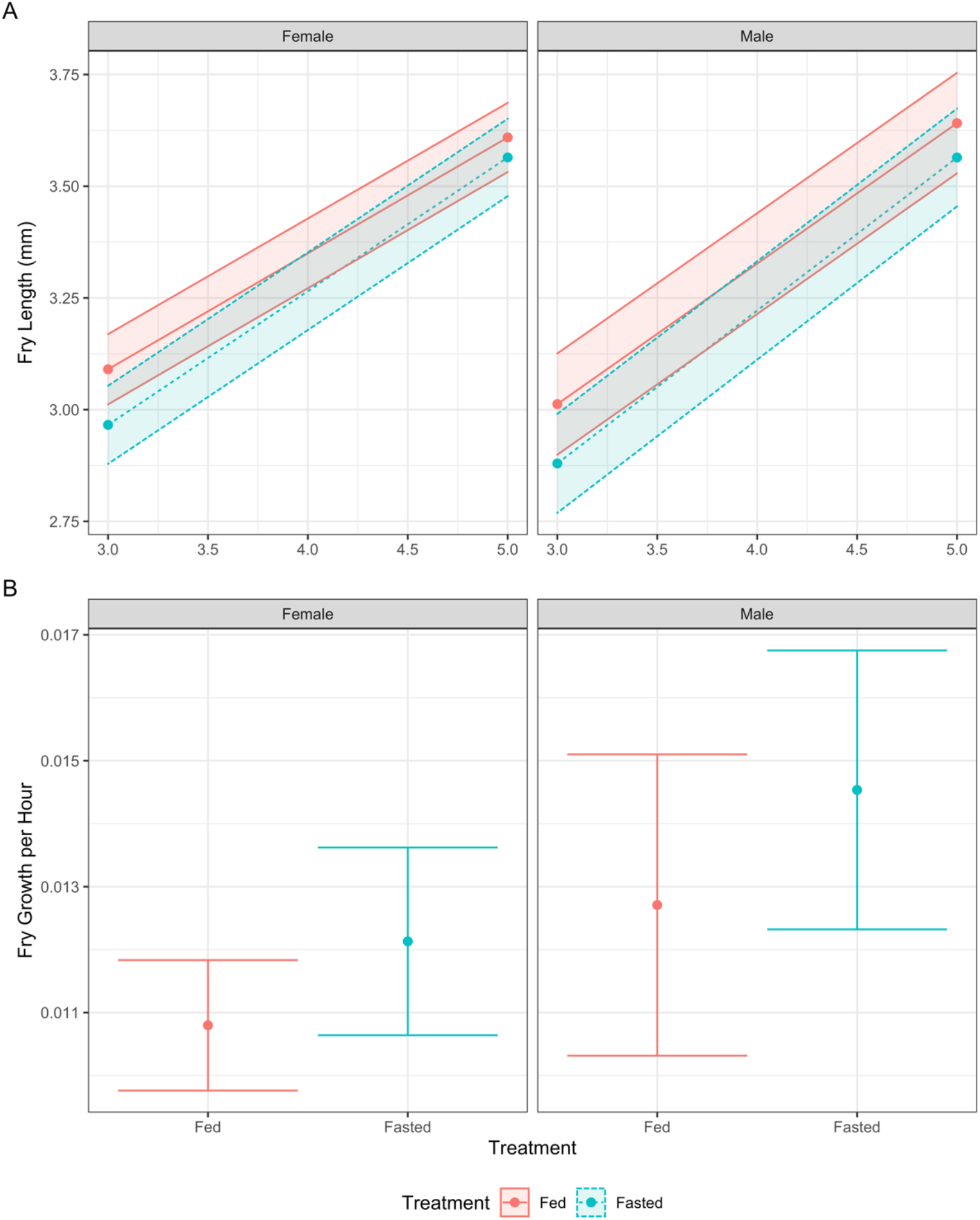
A) Parental sex-specific fry growth between days 3 and 5 postfertilization. Colours represent fasted (blue) or fed (red) dietary treatments in female (left) or male parents (right). B) Hourly fry growth in either female (left, red) or male parents (right, blue). Lines, points and 95% confidence intervals represent predicted values from a linear model.

## Discussion

During periods of dietary restriction, organisms increase investment into somatic maintenance and survival but lower investment into reproduction [5]. However, investment in reproduction can be subdivided into investment in offspring number and offspring quality (via germline maintenance and parental care) [39,40]. The putative trade-offs may comprise different combinations of traits [39,40]. Whilst we detected no DR-related increase in investment into the soma when organisms were on low food, there was evidence of a trade-off between female fecundity, which was negatively affected, and offspring quality, which was maintained at a consistently high level (around ~95%). Prioritising offspring quality over quantity under periods of restricted food is a classic example of a commonly observed life history trade-off [41–43]. Strikingly, however, this trade-off reversed upon refeeding in female zebrafish. Female reproduction increased, which compensated for the period of reduced nutrient intake, but it did so in conjunction with a reduction in offspring survival. A similar refeeding-related increase in reproduction following transient fasting was recently reported in *Drosophila melanogaster* [7].

However, here we show that the increase in female reproduction is associated with a significant cost to gamete quality resulting in reduced offspring survival. Furthermore, the increase in offspring number was associated with significant investment into the soma, denoted with the increased fin regrowth rate upon refeeding. These results further suggest that trade-offs between somatic maintenance, offspring number and offspring quality are dynamic, and organisms change their allocation strategies depending on the environment. Specifically, post-DR trade-offs differ from what is typically observed, with females trading offspring quality for improved somatic maintenance and increased fecundity (Fig. 6). This suggests that whilst investment into germline production was occurring, there were not sufficient resources allocated into germline maintenance [39,40,44,45]. As a result, we could have expected an increase in germline mutations [40,44], which ultimately contributed to the reduced egg quality and egg survival during female refeeding. Indeed, this effect may persist across multiple generations and produce not just inter-but also transgenerational effects of diet, as suggested in a recent study in *Drosophila* [46].

**Figure 6.**
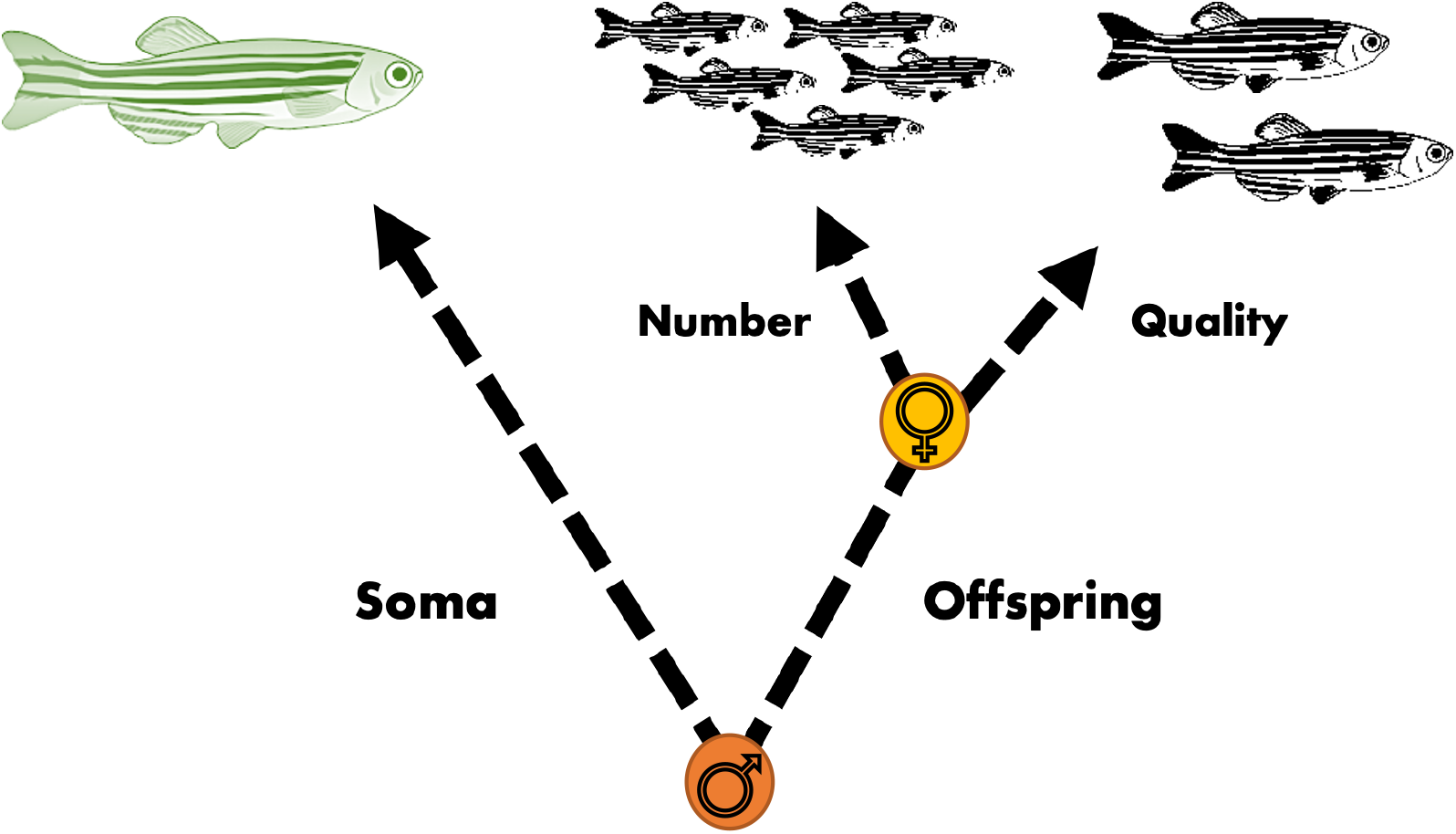
Proposed allocation decisions of females (yellow) and males (orange) after fasting when organisms resume full-feeding. We note that for males, offspring traits are measured using a control female. Females trade offspring quality for offspring number and somatic maintenance. Males trade offspring number and quality for somatic maintenance. The orange or yellow dot denotes where the trade-off was observed during this experiment.

A similar decline in gamete quality was found in males, however our results indicated that whilst a reversible trade-off occurred between offspring quality and quantity in females, no such reverse occurred for males. This could in part be due to the fact that reproduction was measured using a control female. If females compensate for declining male quality, as suggested in a variety of different species [47,48], then reproductive number could be maintained at a high level regardless of the dietary environment of the male. We did, however, detect a significant decline in sperm quality, here measured as sperm VCL, which further increased during the period of refeeding. This coupled with the significant reduction in egg survival both during and after the fasting periods, suggest that regardless of this female compensation, males had traded-off gamete quality with investment into the soma (Fig. 6).

Interestingly, offspring from fasted males exhibited significantly faster growth rate in comparison to those from *ad libitum* males. A similar trend was observed in offspring from the experimental females (albeit not statistically detectable) suggesting a form of compensatory growth. Such increased growth as a result of poor parental conditions has been found in another fish species, *Acanthochromis polyacanthus*. Here, juveniles from low quality parental diets, exhibited a sufficient enough increase in growth to catch up to the length of individuals from high quality parents (regardless of their own dietary environment) [25]. A similar effect was also found in three-spined sticklebacks (*Gasterosteus aculeatus*), however in this particular study the effects were only investigated in the manipulated generation [49]. As a result of this adaptive increase in growth, we may expect costs to manifest in other traits, for instance in the strength of certain skeletal elements [50] or in a reduction in median lifespan [49].

Taken together, the detrimental effects of transient fasting manifested both during and, importantly, after fasting in both male and female gametes, in addition to the potential costs of compensatory offspring growth, suggest that future dietary interventions, particularly those aimed at healthspan extension in humans, should carefully evaluate the impacts of such regimes beyond the parental generation. Our findings show that transient fasting differentially affects gamete quality during and after fasting and that such effects depend on sex in a model vertebrate. Combined with previous work in invertebrate models, this research highlights the urgent need to investigate both short- and longterm effects of different forms of dietary restriction, and other interventions aimed at healthspan extension, on gamete quality and offspring health and fitness.

## Acknowledgements

AAM was funded by ERC and the BBSRC; SI: was funded by ERC and NERC.

The authors would like to thank the staff from the disease monitoring unit, (Simon Deakin, Jonathan Drury, Emma Penwolf, Anya Croft & Imogen Williams) at the University of East Anglia for helping with feeding and maintenance of the general zebrafish stocks from which the samples were selected. We would also like to thank Adrian Rat, Andreas Sutter, Mabel Sydney and Daniel Marcu for helping with data collection and entry.

