## Supplementary materials for "Fasting increases investment in soma upon refeeding at the cost of gamete quality in zebrafish"

Table S1. The effect of fasting on various life history traits either *during* the fasting period or upon *refeeding*. An upwards arrow indicates a significantly positive effect of fasting in comparison to *ad libitum*, whereas a downward arrow indicates the opposite. A dash means no effect was found – or in the case of sperm VCL was not measured in female zebrafish. For reproduction traits (namely, ARS, LRS and egg survival), the effect measured in males is instead the effect of fasted or fed males on a control wildtype female. Results are from (parental) sex-specific models only.

| Trait | ***During Fasting*** | | ***Refeeding*** | | ***Overall*** | |
|---|---|---|---|---|---|---|
| | Male | Female | Male | Female | Male | Female |
| Fin Regrowth | - | - | - | ↑ | | |
| ARS | - | - | - | - | | |
| LRS | - | ↓ | - | - | - | - |
| Egg Survival | ↓ | ↓ | ↓ | ↓ | | |
| Fertility | - | ↓ | - | - | | |

| | | | | |
|---|---|---|---|---|
| Sperm VCL | ↑ | NA | ↓ | NA |
| Fry Growth/Day | - | - | NA | NA |
| Fry Growth/Hour | ↑ | - | NA | NA |

Table S2. Full model output for the three-way mixed model of fin growth per day with both sexes included *during the fasting period.*

| *Predictors* | *Estimates* | *Std. Error* | *Statistic* | *P* |
|---|---|---|---|---|
| (Intercept) | 2.54 | 0.55 | 4.64 | **<0.001** |
| Treatment [Fasted] | -0.84 | 0.59 | -1.43 | 0.152 |
| Sex [Male] | 0.77 | 0.63 | 1.23 | 0.217 |
| Day | 0.37 | 0.04 | 9.33 | **<0.001** |
| Treatment [Fasted] * Sex [Male] | -0.45 | 0.81 | -0.56 | 0.578 |
| Treatment [Fasted] * Day | 0.02 | 0.05 | 0.32 | 0.746 |
| Sex [Male] * Day | -0.04 | 0.06 | -0.68 | 0.495 |
| Treatment [Fasted] * Sex [Male] * Day | 0.08 | 0.07 | 1.13 | 0.260 |
| Observations | 175 | | | |

Table S3. Full model output for the two-way mixed model of fin growth per day with both sexes included *during the refeeding period.*

| *Predictors* | *Estimates* | *Std. Error* | *Statistic* | *P* |
|---|---|---|---|---|
| (Intercept) | 3.75 | 0.81 | 4.64 | **<0.001** |
| Treatment [Fasted] | -1.06 | 0.71 | -1.50 | 0.134 |
| Sex [Male] | 1.56 | 0.72 | 2.17 | **0.030** |
| Day | 0.25 | 0.02 | 14.41 | **<0.001** |

| | | | | |
|---|---|---|---|---|
| Treatment [Fasted] * Sex [Male] | -0.52 | 0.76 | -0.68 | 0.495 |
| Treatment [Fasted] * Day | 0.05 | 0.02 | 2.35 | **0.019** |
| Sex [Male] * Day | -0.06 | 0.02 | -3.21 | **0.001** |
| Observations | 195 | | | |

Table S4. Full model output for the two-way mixed model of fin growth per day for females *during the refeeding period.*

| *Predictors* | *Estimates* | *Std. Error* | *Statistic* | *P* |
|---|---|---|---|---|
| (Intercept) | 6.01 | 1.42 | 4.23 | **<0.001** |
| Treatment [Fasted] | -4.69 | 1.92 | -2.45 | **0.014** |
| Day | 0.16 | 0.04 | 3.99 | **<0.001** |
| Treatment [Fasted] * Day | 0.20 | 0.06 | 3.34 | **0.001** |
| Observations | 69 | | | |

Table S5. Full model output for the one-way mixed model of age-specific reproduction for females *during the fasting period.*

| *Predictors* | *Log-Mean* | *Std. Error* | *Statistic* | *P* |
|---|---|---|---|---|
| (Intercept) | 3.56 | 0.28 | 12.53 | **<0.001** |
| Day | 0.07 | 0.02 | 3.68 | **<0.001** |
| Treatment [Fasted] | -0.65 | 0.26 | -2.46 | **0.014** |
| **Zero-Inflated Model** | | | | |
| (Intercept) | -1.39 | 0.32 | -4.30 | **<0.001** |
| Observations | 60 | | | |

Table S6. Full model output for the one-way mixed model of LRS for females *during the fasting period.*

| *Predictors* | *Log-Mean* | *Std. Error* | *Statistic* | *P* |
|---|---|---|---|---|
| (Intercept) | 4.93 | 0.37 | 13.35 | **<0.001** |

| | | | | |
|---|---|---|---|---|
| Treatment [Fasted] | -0.77 | 0.34 | -2.29 | **0.022** |
| Observations | 30 | | | |

Table S7. Full model output for the one-way mixed model of LRS for males *during the fasting period.*

| *Predictors* | *Log-Mean* | *Std. Error* | *Statistic* | *P* |
|---|---|---|---|---|
| (Intercept) | 5.16 | 0.30 | 17.14 | **<0.001** |
| Treatment [Fasted] | -0.08 | 0.23 | -0.35 | 0.724 |
| **Zero-Inflated Model** | | | | |
| (Intercept) | -2.64 | 0.73 | -3.61 | **<0.001** |
| Observations | 30 | | | |

Table S8. Full model output for the one-way mixed model of age-specific reproduction for females *during the refeeding period.*

| *Predictors* | *Log-Mean* | *Std. Error* | *Statistic* | *P* |
|---|---|---|---|---|
| (Intercept) | 4.39 | 0.42 | 10.41 | **<0.001** |
| Day | 0.00 | 0.01 | 0.21 | 0.833 |
| Treatment [Fasted] | 0.12 | 0.22 | 0.58 | 0.564 |
| **Zero-Inflated Model** | | | | |
| (Intercept) | -1.83 | 0.34 | -5.35 | **<0.001** |
| Observations | 72 | | | |

Table S7. Full model output for the one-way mixed model of LRS for females *during the refeeding period.*

| *Predictors* | *Log-Mean* | *Std. Error* | *Statistic* | *P* |
|---|---|---|---|---|
| (Intercept) | 5.48 | 0.16 | 33.42 | **<0.001** |
| Treatment [Fasted] | 0.16 | 0.23 | 0.67 | 0.502 |
| Observations | 24 | | | |

Table S9. Full model output for the one-way mixed model of LRS for males *during the refeeding period.*

| *Predictors* | *Log-Mean* | *Std. Error* | *Statistic* | *P* |
|---|---|---|---|---|
| (Intercept) | 5.72 | 0.25 | 22.83 | **<0.001** |
| Treatment [Fasted] | -0.33 | 0.23 | -1.45 | 0.148 |
| Observations | 24 | | | |

Table S10. Full model output for the two-way mixed model of egg survival for females *during the fasting period.*

| *Predictors* | *Log-Odds* | *Std. Error* | *Statistic* | *P* |
|---|---|---|---|---|
| (Intercept) | 1.71 | 0.63 | 2.70 | **0.007** |
| Treatment [Fasted] | 2.21 | 0.82 | 2.68 | **0.007** |
| Day | 0.05 | 0.02 | 2.31 | **0.021** |
| Treatment [Fasted] * Day | -0.12 | 0.05 | -2.60 | **0.009** |
| Observations | 3275 | | | |

Table S11. Full model output for the two-way mixed model of egg survival for males *during the fasting period.*

| *Predictors* | *Log-Odds* | *Std. Error* | *Statistic* | *P* |
|---|---|---|---|---|
| (Intercept) | 2.70 | 0.64 | 4.22 | **<0.001** |
| Treatment [Fasted] | 4.87 | 0.88 | 5.54 | **<0.001** |
| Day | -0.01 | 0.02 | -0.55 | 0.581 |
| Treatment [Fasted] * day | -0.46 | 0.03 | -14.96 | **<0.001** |
| Observations | 5102 | | | |

Table S12. Full model output for the two-way mixed model of egg survival for males *during the refeeding period.*

| *Predictors* | *Log-Odds* | *Std. Error* | *Statistic* | *P* |
|---|---|---|---|---|
| (Intercept) | 1.92 | 0.71 | 2.69 | **0.007** |
| Treatment [Fasted] | 3.96 | 1.14 | 3.48 | **0.001** |
| Day | 0.02 | 0.01 | 1.68 | 0.092 |

| | | | | |
|---|---|---|---|---|
| Treatment [Fasted] * Day | -0.16 | 0.02 | -10.35 | **<0.001** |
| Observations | 7229 | | | |

Table S13. Full model output for the two-way mixed model of egg survival for females *during the refeeding period.*

| *Predictors* | *Log-Odds* | *Std. Error* | *Statistic* | *P* |
|---|---|---|---|---|
| (Intercept) | -0.13 | 0.63 | -0.21 | 0.833 |
| Treatment [Fasted] | 5.68 | 0.89 | 6.36 | **<0.001** |
| Day | 0.11 | 0.01 | 7.74 | **<0.001** |
| Treatment [Fasted] * Day | -0.25 | 0.02 | -13.38 | **<0.001** |
| Observations | 6249 | | | |

Table S14. Full model output for the two-way mixed model of sperm VCL *during the fasting period.*

| *Predictors* | *Estimates* | *Std. Error* | *Statistic* | *P* |
|---|---|---|---|---|
| (Intercept) | 4.23 | 0.03 | 122.81 | **<0.001** |
| Treatment [Fasted] | -0.06 | 0.04 | -1.38 | 0.168 |
| Day | -0.01 | 0.00 | -9.68 | **<0.001** |
| Replicate Block [B2] | -0.04 | 0.04 | -1.05 | 0.294 |
| Treatment [Fasted] * Day | 0.01 | 0.00 | 3.87 | **<0.001** |
| Observations | 67449 | | | |

Table S15. Full model output for the two-way mixed model of sperm VCL *during the refeeding period.*

| *Predictors* | *Estimates* | *Std. Error* | *Statistic* | *P* |
|---|---|---|---|---|
| (Intercept) | 4.28 | 0.05 | 79.97 | **<0.001** |
| Treatment [Fasted] | 0.19 | 0.07 | 2.81 | **0.005** |
| Day | -0.01 | 0.00 | -11.55 | **<0.001** |
| Replicate Block [B2] | 0.06 | 0.06 | 1.06 | 0.291 |
| Treatment [Fasted] * Day | -0.01 | 0.00 | -7.92 | **<0.001** |
| Observations | 25229 | | | |

Table S16. Full model output for the two-way mixed model of fry growth per day.

| *Predictors* | *Estimates* | *Std. Error* | *Statistic* | *P* |
|---|---|---|---|---|
| (Intercept) | 2.24 | 0.07 | 30.06 | **<0.001** |
| Treatment [Fasted] | -0.21 | 0.06 | -3.31 | **0.001** |
| Day | 0.26 | 0.01 | 21.81 | **<0.001** |
| Sex [Male] | -0.17 | 0.06 | -2.62 | **0.009** |
| Treatment [Fasted] * Day | 0.03 | 0.01 | 2.24 | **0.025** |
| Treatment [Fasted] * Sex [Male] | -0.02 | 0.03 | -0.71 | 0.479 |
| Day * Sex [Male] | 0.05 | 0.01 | 3.49 | **<0.001** |
| Observations | 1728 | | | |

Table S17. Full model output for the two-way mixed model of fry growth per day from experimental females.

| *Predictors* | *Estimates* | *Std. Error* | *Statistic* | *P* |
|---|---|---|---|---|
| (Intercept) | 2.31 | 0.06 | 36.49 | **<0.001** |
| Treatment [Fasted] | -0.24 | 0.09 | -2.63 | **0.008** |
| Day | 0.26 | 0.01 | 20.78 | **<0.001** |
| Treatment [Fasted] * Day | 0.04 | 0.02 | 1.79 | 0.073 |
| Observations | 709 | | | |

Table S18. Full model output for the two-way mixed model of fry growth per day from experimental males.

| *Predictors* | *Estimates* | *Std. Error* | *Statistic* | *P* |
|---|---|---|---|---|
| (Intercept) | 2.07 | 0.08 | 26.08 | **<0.001** |
| Treatment [Fasted] | -0.22 | 0.08 | -2.67 | **0.008** |
| Day | 0.31 | 0.01 | 22.63 | **<0.001** |
| Treatment [Fasted] * Day | 0.03 | 0.02 | 1.45 | 0.148 |

| | |
|---|---|
| Observations | 1019 |

Table S19. Full model output for the one-way model of average hourly fry growth rate across both parental sexes.

| *Predictors* | *Estimates* | *Std. Error* | *Statistic* | *P* |
|---|---|---|---|---|
| (Intercept) | 0.01005 | 0.00073 | 13.82 | **<0.001** |
| Treatment [Fasted] | 0.00158 | 0.00061 | 2.57 | **0.010** |
| Sex [Male] | 0.00187 | 0.00061 | 3.05 | **0.002** |
| Replicate Block [2] | 0.00015 | 0.00083 | 0.18 | 0.854 |
| Replicate Block [3] | -0.00076 | 0.00081 | -0.93 | 0.350 |
| Replicate Block [4] | 0.00405 | 0.00091 | 4.45 | **<0.001** |
| Replicate Block [5a] | -0.00128 | 0.00180 | -0.71 | 0.479 |
| Replicate Block [5b] | 0.00351 | 0.00590 | 0.59 | 0.552 |
| Observations | 802 | | | |

Table S20. Full model output for the one-way model of average hourly fry growth rate from experimental males.

| *Predictors* | *Estimates* | *Std. Error* | *Statistic* | *P* |
|---|---|---|---|---|
| (Intercept) | 0.01261 | 0.00086 | 14.66 | **<0.001** |
| Treatment [Fasted] | 0.00183 | 0.00083 | 2.20 | **0.028** |
| Replicate Block [2] | -0.00041 | 0.00111 | -0.37 | 0.713 |
| Replicate Block [3] | -0.00288 | 0.00113 | -2.55 | **0.011** |
| Replicate Block [4] | 0.00352 | 0.00122 | 2.89 | **0.004** |
| Replicate Block [5a] | -0.00221 | 0.00195 | -1.13 | 0.257 |
| Replicate Block [5b] | 0.00257 | 0.00616 | 0.42 | 0.676 |
| Observations | 469 | | | |

Table S21. Full model output for the one-way model of average hourly fry growth rate from experimental females.

| *Predictors* | *Estimates* | *Std. Error* | *Statistic* | *P* |
|---|---|---|---|---|
| (Intercept) | 0.00880 | 0.00096 | 9.17 | **<0.001** |
| Treatment [Fasted] | 0.00133 | 0.00092 | 1.46 | 0.145 |
| Replicate Block [2] | 0.00109 | 0.00124 | 0.88 | 0.380 |
| Replicate Block [3] | 0.00187 | 0.00116 | 1.61 | 0.107 |
| Replicate Block [4] | 0.00505 | 0.00136 | 3.73 | **<0.001** |
| Observations | 333 | | | |
